# Brain pericyte lineages converge on a shared molecular profile

**DOI:** 10.64898/2026.11.24.768980

**Authors:** Aidan F. Durkan, Jacklyn Levey, Mallika Shankar, Natasha M. O’Brown

## Abstract

Brain pericytes arise from two embryonic lineages, neural crest and mesoderm, but it is unknown whether these origins produce molecularly distinct pericyte populations. Using zebrafish lineage-tracing transgenics, we mapped neural crest- and mesoderm-derived pericytes across the brain from larval to adult stages. Neural crest-derived pericytes occupied the forebrain and mesoderm-derived pericytes occupied the midbrain and hindbrain at early stages. A population of neural crest-derived pericytes then appeared in the dorsal midbrain by 10 days post-fertilization and persisted into adulthood. A small population of mesoderm-derived pericytes also appeared in the ventral forebrain of juvenile and adult fish. Bulk RNA sequencing of FACS-isolated adult pericytes showed that the two lineages share expression of canonical pericyte genes, with only 30 genes differentially expressed. Fluorescent *in situ* hybridization validated one neural crest-enriched gene, *plpp3*, and several mesoderm-enriched genes, including *loxl2b*. Together, these data suggest that despite distinct embryonic origins and regional bias, brain pericyte lineages converge on a largely shared molecular profile.

## INTRODUCTION

Mural cells, including vascular smooth muscle cells and pericytes, surround the endothelium of blood vessels and support vessel stability^1^. In the central nervous system (CNS), pericytes are enriched on capillaries, where they are embedded in the endothelial basement membrane and form specialized contacts that are critical for microvascular integrity^2,3^. Across vertebrates, mural cells are commonly identified using a conserved panel of markers, including *pdgfrb* and *notch3*^4^, which are broadly expressed by both pericytes and vascular smooth muscle cells, and *tagln* and *acta2*, which mark vascular smooth muscle cells^4,5^. In zebrafish, mural cells are visible on the cerebral vasculature at 48 hours post-fertilization (hpf), based on *pdgfrb* and *notch3 in situ* hybridization^4^; *pdgfrb*+ cells are also detected at 48 hpf on the basilar artery and primordial hindbrain channels, and by 60 hpf they begin extending onto the cerebral central arteries, where their numbers steadily increase^6^. By 120 hpf, *pdgfrb*+ cells extensively cover the central cerebral arteries throughout the brain^6^.

Classic quail–chick chimera studies demonstrated that brain mural cells arise from two discrete embryonic lineages: neural crest and mesoderm^7-9^. These early lineage tracing studies indicated that the neural crest contributes substantially to pericytes in the forebrain and the mesoderm to those in the midbrain and hindbrain. Consistent with a neural crest contribution, knockdown of *foxd3* and /or *tfap2a*, which are required for early neural crest specification, reduces the number of mural cells in the forebrain^4,5,10^. Conversely, knockdown of *tbx6*, a gene required for early mesoderm specification, results in a decrease in hindbrain pericytes^6^. These findings suggested regional differences in pericyte lineage, with neural crest-derived pericytes predominating in the forebrain and mesoderm-derived pericytes in the hindbrain. However, while these experiments established discrete developmental origins for CNS pericytes, it remains unresolved whether these different lineages produce molecularly distinct pericyte populations.

Pericytes are required for blood–brain barrier (BBB) development and function. Mouse studies have demonstrated that this dependence is mediated largely through PDGFB–PDGFRβ signaling^2,3,11^. Endothelial cells secrete the ligand PDGFB, which signals to PDGFRβ on mural cells, driving their recruitment to and proliferation along the vessel wall^2,3,11^. Perturbations to this pathway disrupt pericyte coverage across vertebrate species^2,3,11-13^. In mice, hypomorphic *Pdgfb* mutants show a dose-dependent relationship in which the degree of pericyte loss correlates with the magnitude of BBB leakiness, accompanied by endothelial transcriptional changes including increased expression of permeability-associated genes such as *Angpt2* and *Plvap*^14^. Mechanistically, this increased permeability results from elevated endothelial transcytosis and altered tight junction organization^2,3,14,15^. Beyond development, PDGFB–PDGFRβ signaling is continuously required to maintain the BBB: inducible, endothelial-specific ablation of Pdgfb in adult mice causes progressive pericyte depletion followed by increased BBB permeability, without other structural vascular defects^16^. Consistent with these findings, chronic pericyte loss in adult mice is associated with capillary rarefaction, hypoperfusion, BBB breakdown, and neurodegeneration^15^. Importantly, these models perturb mural cell coverage broadly and do not resolve whether all pericyte types contribute equally to BBB integrity, or whether specific pericyte subsets, such as those of distinct developmental origins, differentially regulate BBB function.

To investigate how developmental origin shapes pericyte identity, we used zebrafish lineage-tracing lines^6^ to map neural crest- and mesoderm-derived pericytes throughout the brain from larval stages to adulthood. To test whether these spatially segregated lineages also differ at the molecular level, we performed RNA sequencing on adult pericytes from both lineages, which revealed markedly similar transcriptomic profiles, with very few differentially expressed genes. Overall, these results suggest that despite their distinct embryonic origins and regional stratification, the two pericyte lineages largely converge on a common molecular and functional state within the mature neurovascular unit.

## RESULTS

### Neural crest- and mesoderm-derived pericytes primarily populate discrete brain regions

To determine where and when different pericyte populations enter the brain, we used previously generated lineage-tracing transgenic lines^6^. Specifically, we used *sox10:Cre; pdgfrb:Gal4ff; UAS:lox-mCherry-lox-mVenus* triple transgenic fish, in which mesoderm-derived pericytes are labeled by mCherry and neural crest-derived pericytes by mVenus (Figure 1). At 3 days post-fertilization (dpf), the two populations were already clearly separated: neural crest-derived pericytes were found exclusively in the forebrain, and mesoderm-derived pericytes in the midbrain and hindbrain (Figure 1A, B). This regional pattern was largely maintained from 3 to 7 dpf (Figure 1A, B). From 10 dpf onward, however, a small population of neural crest-derived pericytes appeared in the dorsal midbrain in 43% of fish (N = 7), indicated by white arrows, and by 1 month all fish had such a population (N=3; Figure 1B, C). This population was largely restricted to one midbrain hemisphere and persisted into adulthood (Figure 1B, C). Its extent varied between fish, and the population was usually concentrated in one midbrain hemisphere, with no consistent side. Neural crest-derived pericytes in the midbrain also appeared denser dorsally and sparser ventrally. We also found that choroid plexus pericytes were exclusively neural crest-derived in the 14-month-old adult fish (Figure 1D). In addition, adult fish had a small population of mesoderm-derived pericytes in the ventral forebrain, at the forebrain-midbrain boundary (Figure 1C, D). Together, these data show that neural crest- and mesoderm-derived pericytes occupy largely distinct brain regions early in development. Still, their territories become partially intermixed at the forebrain-midbrain boundary as the animal matures.

**Figure 1.**
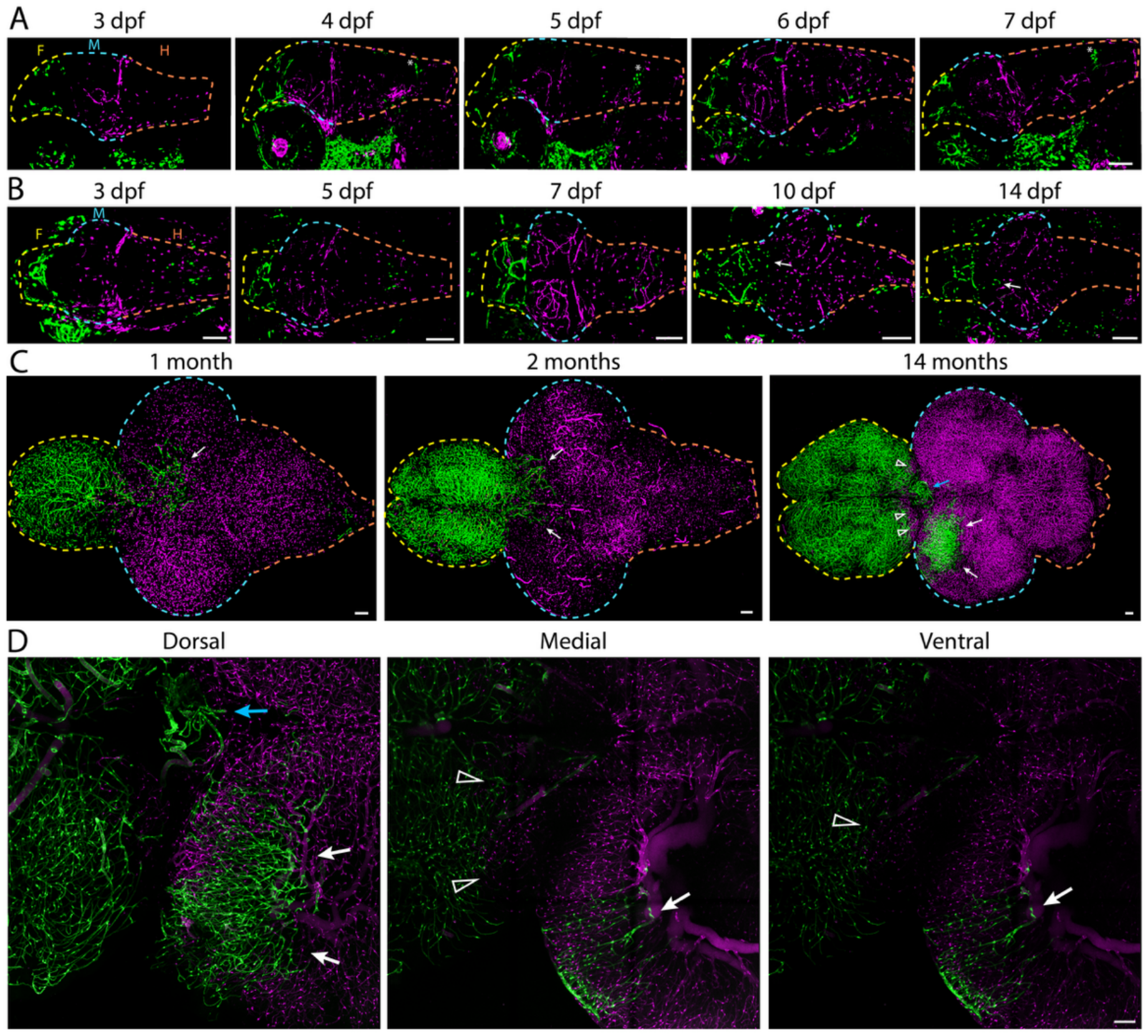
Lineage tracing across the lifespan reveals discrete locations for different pericyte populations. **A**. Sagittal views of *sox10:Cre; pdgfrb:Gal4ff; UAS:lox-mCherry-lox-mVenus* triple transgenic fish reveal neural crest-derived pericytes (green) exclusively in the forebrain (F, yellow dashed line) from 3 to 7 dpf (3 dpf, n = 10; 4 dpf, n = 5; 5 dpf, n = 8; 6 dpf, n = 11; 7 dpf, n = 7), and mesoderm-derived pericytes (magenta) in the midbrain (M, blue dashed line) and hindbrain (H, orange dashed line). A population of neural crest-derived neurons (*) in the hindbrain is also labeled in these transgenics. **B**. Dorsal views of triple transgenic fish reveal some neural crest-derived pericytes entering the midbrain at 10 dpf (n = 7) and 14 dpf (n = 4) (white arrows). **C**. CUBIC-cleared brains from juvenile (1 month, n = 3; 2 months, n = 2) and adult (14 months, n = 2) fish show that the midbrain neural crest-derived population is maintained and expands with age (white arrows). Pericytes in the choroid plexus (blue arrow) are exclusively neural crest-derived. Mesoderm-derived pericytes are also present in the posterior forebrain of the 14-month-old fish (arrowheads). **D**. Magnified maximum intensity projections (300 µm thick) spanning the midbrain-hindbrain boundary of the 14-month-old fish show neural crest-derived pericytes in the midbrain (white arrows) and in the choroid plexus (blue arrow), with mesoderm-derived pericytes in the medial and ventral regions of the posterior hindbrain (arrowheads). Scale bars are 100 µm.

### Neural crest and mesoderm-derived pericyte transcriptomics are largely similar

To determine whether pericyte origin affects molecular identity, we performed bulk RNA-sequencing on fluorescence-activated cell (FAC)-sorted neural crest- and mesoderm-derived pericytes from adult triple transgenic brains (Figure 2A). Differential gene expression analysis showed that most genes were expressed at similar levels in the two populations (Supplemental Table 1), including all canonical pericyte genes, such as *kcnj8, ednraa, foxc1b, mcamb, ndufa4l2a, notch3*, and *pdgfrb* (Figure 2B). Only 30 genes were significantly differentially expressed, 19 enriched in neural crest-derived pericytes and 11 in mesoderm-derived pericytes (Figure 2B). To validate these differences, we performed hybridization chain reaction fluorescent *in situ* hybridization (HCR-FISH) in adult animals for the differentially expressed genes with expression of at least 20 FPKM in our RNA-seq data. Among the neural crest-enriched genes, only *plpp3* was confirmed by HCR-FISH, whereas *tfap2a* showed significantly higher signal in mesoderm-derived pericytes, opposite to the RNA-seq result (Figure 2C, F). In contrast, several mesoderm-enriched genes were confirmed, including *loxl2b, plin2, cx32*.*2, lgals9l1*, and *ccl35*.*1* (Figure 2D, F), with the strongest effect seen for *loxl2b*. Consistent with the RNA-seq data, we saw no significant difference in the general pericyte markers *notch3* and *pdgfrb* (Figure 2E, F).

**Figure 2.**
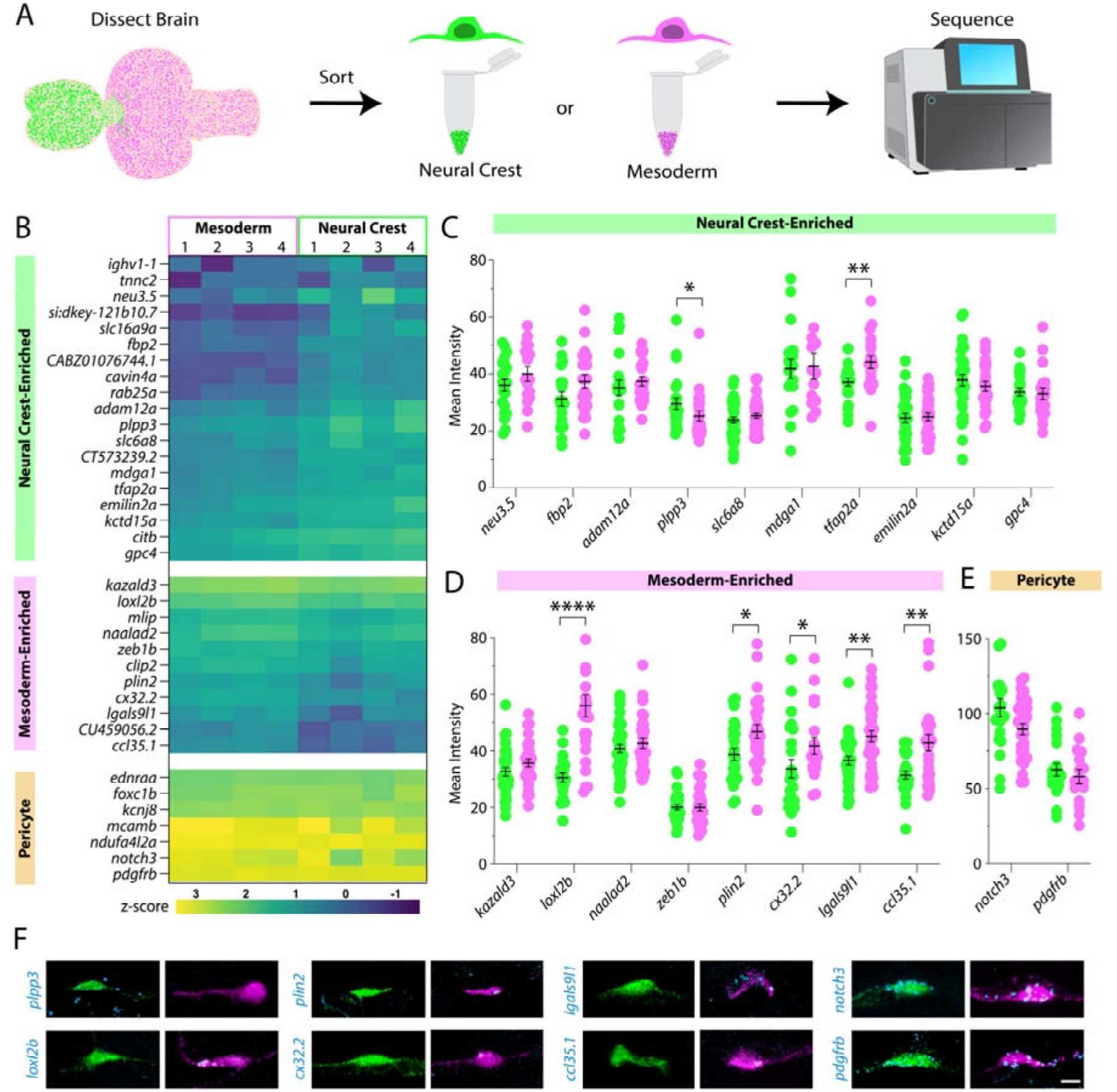
Pericyte origin has limited effects on gene expression. **A**. Schematic of RNA-seq library preparation from FACS-sorted neural crest-derived (green) and mesoderm-derived (magenta) pericytes from dissected adult zebrafish brains. **B**. Heatmap of the 30 differentially expressed genes (19 neural crest-enriched, 11 mesoderm-enriched) and seven canonical pericyte genes across four biological replicates per population. **C-E**. HCR-FISH quantification of mean transcript signal intensity in neural crest-derived (green) and mesoderm-derived (magenta) pericytes for differentially expressed genes with expression of at least 20 FPKM. Of the neural crest-enriched genes (C), *plpp3* is significantly higher in neural crest-derived pericytes, whereas *tfap2a* is significantly higher in mesoderm-derived pericytes. Among mesoderm-enriched genes (D), *loxl2b, plin2, cx32*.*2, lgals9l1*, and *ccl35*.*1* are significantly higher in mesoderm-derived pericytes. The general pericyte genes *notch3* and *pdgfrb* (E) show no difference. Each point represents a single pericyte; bars show mean ± SEM. **F**. Representative images of neural crest-derived (left, green) and mesoderm-derived (right, magenta) pericytes with HCR signal for the indicated transcript (blue). * p < 0.05, ** p < 0.01, **** p < 0.0001 by multiple Mann-Whitney tests. Scale bar is 5 µm.

## DISCUSSION

Brain pericytes arise from two distinct embryonic lineages, and the long-standing model holds that neural crest-derived pericytes are restricted to the forebrain. In contrast, mesoderm-derived pericytes populate the midbrain, hindbrain, and remaining CNS^6,8,17-19^. This model stems from early quail–chick chimera studies, which showed that neural crest-derived cells heavily populate the forebrain while mesodermal cells give rise to pericytes throughout the rest of the CNS ^8,17^. It was later corroborated by lineage-tracing studies in mice, which demonstrated that cephalic neural crest-derived cells migrate into the developing telencephalon around E9.5 and differentiate into pericytes (Pdgfrβ+ /NG2+) by E11.5, while mesoderm-derived pericytes are largely excluded from this region^18^. Most existing zebrafish data are consistent with this pattern, with neural crest-derived pericytes populating the forebrain and mesoderm-derived pericytes populating the midbrain and hindbrain^6,19^. However, the distribution of these two lineages across brain regions has not been mapped in detail from larval development through adulthood, and whether their origin drives differences in molecular identity in the mature brain has not been systematically examined.

Using previously established pericyte lineage-tracing fish^10^, we found that early larval patterns agree with this model, but the regional separation begins to break down at later larval stages (Figure 1B). These data refine the long-standing model, showing that lineage territories are regionally biased but not strictly fixed, with the lineages intermixing at the forebrain-midbrain boundary into adulthood (Figure 1C-D). Despite these spatial differences in origin, adult pericytes from both lineages expressed canonical pericyte genes at similar levels, with only 30 genes differentially expressed overall (Figure 2; Supplementary Table 1). One candidate route to this convergence is a shared *nkx3*.*1*-expressing, fibroblast-like precursor state identified by Ahuja et al. in pericyte development from both germ layers; however, because we profiled only adult animals, our data capture similarity at maturity rather than a trajectory of convergence over time^20^. The genes that did differ, *plpp3* in neural crest-derived pericytes and l*oxl2b, plin2, cx32*.*2, lgals9l1*, and *ccl35*.*1* in mesoderm-derived pericytes, differed modestly, with low transcript abundance and signal in both lineages, so they are best described as lineage-enriched rather than lineage-exclusive. *Tfap2a* was neural crest-enriched by RNA-seq but higher in mesoderm-derived pericytes by HCR-FISH, which measured transcript levels in individual pericytes. This discrepancy may reflect low expression levels, which would make both assays noisy, and label non-pericyte cells in the sorted neural crest population, such as the neural crest-derived hindbrain neurons visible in Figure 1. Notably, we also observed that pericytes within the choroid plexus were exclusively neural crest-derived in adulthood (Figure 1C-D), suggesting this structure may represent a site where lineage segregation is strictly maintained despite the broader intermingling seen elsewhere in the brain.

Overall, our analysis shows that neural crest- and mesoderm-derived pericytes are highly similar to one another at the level of gene expression, with lineage playing a very minor role in shaping their transcriptional identity. Our lineage tracing relies exclusively on *sox10:Cre* to mark neural crest-derived pericytes, with the mesoderm-derived population inferred as the unrecombined remainder rather than labeled with a complementary mesoderm-driven *tbx6:Cre* as originally done ^6^. Therefore, our mesoderm population may include a small number of neural crest-derived cells rather than being purely mesoderm-derived. Additionally, because we used bulk RNA sequencing of FACS-sorted cells, our approach was not designed to resolve finer cellular subtypes or to distinguish pericytes from other mural cells, such as vascular smooth muscle cells. Building on these findings, single-cell profiling paired with lineage-selective manipulation will be a valuable next step to further resolve cellular heterogeneity within each lineage and test whether the handful of lineage-enriched genes identified here have functional significance.

## MATERIALS AND METHODS

### Zebrafish Lines and Maintenance

Zebrafish were maintained at 28.5°C following standard protocols ^21^. All work was approved by the Rutgers University Standing Committee under protocol number 202300038. Adult fish were maintained on a standard 12-hour light-dark cycle. Adult fish, aged 3 months to 1.5 years, were crossed to produce embryos and larvae. For larvae used in confocal microscopy, 0.003% phenylthiourea (PTU; Sigma P7629) was applied from 1 dpf to 7 dpf to inhibit melanin production. These studies used the Tg(*sox10:Cre,cryaa:DsRed*)^zy70Tg^ line^22^ abbreviated as *sox10:Cre* in the text, the TgBAC(pdgfrb:GAL4FF)^ncv24Tg^ line^6^ abbreviated as *pdgfrb:Gal4ff* in the text, and the Tg(*5xUAS:LOXP-mCherry-LOXP-mVenus*)^ncv28Tg^ line^6^ abbreviated as *UAS:lox-mCherry-lox-mVenus* in the text.

### Imaging of lineage tracing transgenic fish

All imaging was carried out on *Tg(sox10:Cre; pdgfrb:Gal4ff; UAS:lox-mCherry-lox-mVenus)* triple transgenic fish. Larvae (3 to 14 dpf) were imaged live after anesthesia with tricaine (4 mg / mL;(Pentair TRS1) and embedded in 2% low-melting-point agarose (Promega V2111), then submerged in tricaine-containing embryo water. Juvenile and adult fish were fixed overnight in 4% paraformaldehyde with their brains exposed. Dissected brains were then cleared with CUBIC ^23^ following a similar protocol. Briefly, brains were incubated in 50% CUBIC-L /50% PBST for 1 hour at 37°C with rocking, then transferred to 100% CUBIC-L for at least 4 hours. Cleared brains were mounted in low-gelling-temperature agarose with CUBIC-R. All imaging was performed on a Leica Stellaris DMi8 confocal microscope with a 10x objective (0.75x zoom), a 25x water objective (0.75x zoom for Figure 1D), and 2 µm z-steps. We then used Ilastik (v1.4.2b8)^24^ to segment neural crest- and mesoderm-derived pericyte populations and to remove background signal from the skin.

### Pericyte isolation, RNA extraction and library preparation

Brains were dissected from adult (6–10-month-old) *sox10:Cre; pdgfrb:Gal4ff; UAS:lox-mCherry-lox-mVenus* triple transgenic zebrafish and dissociated into single-cell suspensions using papain. Neural crest–derived and mesoderm-derived cell populations were isolated by fluorescence-activated cell sorting on a FACS Aria 594 (BD Biosciences) based on fluorescent reporter expression. Neural crest–derived cells were collected as mVenus□ events, whereas mesoderm-derived pericytes were collected as mCherry□ events, and sorted cells were deposited directly into TRIzol (Thermo Fisher Scientific) for RNA extraction. Total RNA was extracted following the manufacturer’s instructions for TRIzol and subsequently purified using the QIAGEN MinElute Cleanup Kit. Sequencing libraries were generated from purified RNA using the TAKARA SMARTer Stranded Total RNA-seq Kit v2, optimized for low-input, stranded total RNA profiling on Illumina platforms, and prepared according to the manufacturer’s protocol.

### Sequencing, alignment, and differential expression

Libraries were sequenced on an Illumina NextSeq 1000 using a P2 flow cell to generate 100 bp paired-end reads. Reads were aligned to the zebrafish reference genome (GRCz11) using the splice-aware STAR aligner^25^, and gene-level counts were obtained for downstream analysis. Differential gene expression was assessed in R using DESeq2^26^. Raw sequencing data and gene-level counts have been deposited in the NCBI Gene Expression Omnibus (GEO) under accession number GSE299235.

### Hybridization Chain Reaction (HCR) fluorescent in situ hybridization

HCR fluorescent in situ hybridization was performed on 14-μm-thick cryosections of fixed adult brains with transgenically labeled pericytes as previously described ^27,28^. Briefly, sections were air-dried and re-fixed in 4% paraformaldehyde (PFA) in PBS for 10 min at room temperature. Following fixation, slides were washed in PBS and then permeabilized using 1 μg /ml Proteinase K (ThermoFisher) for 5 min, followed by PBS washes and refixation. Tissues were further permeabilized by an ethanol dehydration series of 50%, 70%, and two rounds of 100% ethanol and air-dried for 5 min. Dried slides were put into probe hybridization solution for at least 10 min at 37°C. Subsequently, probes for different pericyte-enriched genes (Supplemental Table 2, IDT DNA oPools) were added to the slides at a final concentration of 4 nM and hybridized overnight at 37°C. The next day, slides were washed with a series of wash buffers to 5x SSCT (5x SSC with 0.1% Tween 20). Excess liquid was then removed, and samples were immersed in amplification buffer at room temperature for 30 min prior to hairpin amplification (Molecular Instruments), which occurred overnight at room temperature. Slides were then washed in 5x SSCT, then 5x SSC, and mounted with 50% glycerol in PBS containing 0.1% sodium azide.

### HCR Imaging and Quantification

All imaging was performed on a Leica Stellaris DMi8 confocal microscope with a 25x water-immersion objective and 0.5 µm z-steps. We generated a 10 µm-thick maximum-intensity projection centered on the middle of the section using Fiji, and the average probe fluorescence intensity was measured in individual pericyte ROIs. Mean fluorescent intensity for all probes was plotted and analyzed by Multiple Mann-Whitney tests in GraphPad Prism. Zoomed-in representative images were acquired using a 63x oil-immersion objective with 0.3 µm z-steps. We collected tissue autofluorescence in the 405 channel and subtracted that from each channel using Fiji.

## ACKNOWLEDGEMENTS

We thank members of the O’Brown lab for data discussions and comments on the manuscript and the Aquatic Core facility, specifically Katie Flaherty at Rutgers for fish maintenance and care.

## AUTHOR CONTRIBUTIONS

A.F.D. and N.M.O. conceived the project and designed the experiments. A.F.D., J.L., M.S., and N.M.O. conducted the animal experiments and analyzed the data. A.F.D., J.L., and N.M.O. wrote the manuscript.

## FUNDING

This work was supported by the National Institutes of Health R00HD103911 (N.M.O. and A.D. and Howard Hughes Medical Institute (N.M.O., J.L.). N.M.O. is a Howard Hughes Medical Institute Freeman Hrabowski Scholar.

## CONFLICTS OF INTEREST

The authors declare no competing interests.

## DATA AND CODE AVAILABILITY

Bulk RNA-seq data were deposited into the Gene Expression Omnibus database under accession number GSE299235 and are available at the following URL: https://identifiers.org/geo:GSE299235. Differential expression results are provided in Supplemental Table 1.

## Notes

### Competing Interest Statement

The authors have declared no competing interest.

## REFERENCES

1. Gaengel, K., Genove, G., Armulik, A. & Betsholtz, C. Endothelial-mural cell signaling in vascular development and angiogenesis. Arterioscler Thromb Vasc Biol 29, 630–638 (2009). 10.1161/ATVBAHA.107.161521

2. Armulik, A. et al. Pericytes regulate the blood-brain barrier. Nature 468, 557–561 (2010). 10.1038/nature09522

3. Daneman, R., Zhou, L., Kebede, A. A. & Barres, B. A. Pericytes are required for blood-brain barrier integrity during embryogenesis. Nature 468, 562–566 (2010). 10.1038/nature09513

4. Wang, Y., Pan, L., Moens, C. B. & Appel, B. Notch3 establishes brain vascular integrity by regulating pericyte number. Development 141, 307–317 (2014). 10.1242/dev.096107

5. Whitesell, T. R. et al. An α-Smooth Muscle Actin (acta2/αsma) Zebrafish Transgenic Line Marking Vascular Mural Cells and Visceral Smooth Muscle Cells. PLoS ONE 9 (2014). 10.1371/journal.pone.0090590

6. Ando, K. et al. Clarification of mural cell coverage of vascular endothelial cells by live imaging of zebrafish. Development 143, 1328–1339 (2016). 10.1242/dev.132654

7. Le Lievre, C. S. & Le Douarin, N. M. Mesenchymal derivatives of the neural crest: analysis of chimaeric quail and chick embryos. J Embryol Exp Morphol 34, 125–154 (1975).

8. Etchevers, H. C., Vincent, C., Le Douarin, N. M. & Couly, G. F. The cephalic neural crest provides pericytes and smooth muscle cells to all blood vessels of the face and forebrain. Development 128, 1059–1068 (2001). 10.1242/dev.128.7.1059

9. Bergwerff, M., Verberne, M. E., DeRuiter, M. C., Poelmann, R. E. & Gittenberger-de-Groot, A. C. Neural Crest Cell Contribution to the Developing Circulatory System. Circulation Research 82, 221–231 (1998). 10.1161/01.Res.82.2.221

10. Wang, W.-D., Melville, D. B., Montero-Balaguer, M., Hatzopoulos, A. K. & Knapik, E. W. Tfap2a and Foxd3 regulate early steps in the development of the neural crest progenitor population. Developmental Biology 360, 173–185 (2011). 10.1016/j.ydbio.2011.09.019

11. Lindblom, P. et al. Endothelial PDGF-B retention is required for proper investment of pericytes in the microvessel wall. Genes & Development 17, 1835–1840 (2003). 10.1101/gad.266803

12. Ando, K. et al. Conserved and context-dependent roles for pdgfrb signaling during zebrafish vascular mural cell development. Developmental Biology 479, 11–22 (2021). 10.1016/j.ydbio.2021.06.010

13. Baltaci, O. F. et al. Mural cells protect the adult brain from hemorrhage but do not control the blood–brain barrier in developing zebrafish. eLife 14 (2026). 10.7554/eLife.104061

14. Mäe, M. A. et al. Single-Cell Analysis of Blood-Brain Barrier Response to Pericyte Loss. Circulation Research 128 (2021). 10.1161/circresaha.120.317473

15. Bell, R. D. et al. Pericytes Control Key Neurovascular Functions and Neuronal Phenotype in the Adult Brain and during Brain Aging. Neuron 68, 409–427 (2010). 10.1016/j.neuron.2010.09.043

16. Vazquez-Liebanas, E. et al. Adult-induced genetic ablation distinguishes PDGFB roles in blood-brain barrier maintenance and development. Journal of Cerebral Blood Flow & Metabolism 42, 264–279 (2021). 10.1177/0271678×211056395

17. Korn, J., Christ, B. & Kurz, H. Neuroectodermal origin of brain pericytes and vascular smooth muscle cells. J Comp Neurol 442, 78–88 (2002). 10.1002/cne.1423

18. Yamanishi, E., Takahashi, M., Saga, Y. & Osumi, N. Penetration and differentiation of cephalic neural crest-derived cells in the developing mouse telencephalon. Dev Growth Differ 54, 785–800 (2012). 10.1111/dgd.12007

19. Girolamo, F. et al. Neural crest cell-derived pericytes act as pro-angiogenic cells in human neocortex development and gliomas. Fluids Barriers CNS 18, 14 (2021). 10.1186/s12987-021-00242-7

20. Ahuja, S. et al. The development of brain pericytes requires expression of the transcription factor nkx3.1 in intermediate precursors. PLoS Biol 22, e3002590 (2024). 10.1371/journal.pbio.3002590

21. Westerfield, M. he Zebrafish Book; A Guide for the Laboratory Use of Zebrafish (Brachydanio rerio). University of Oregon Press, 300P (1993).

22. Abdul-Wajid, S., Demarest, B. L. & Yost, H. J. Loss of embryonic neural crest derived cardiomyocytes causes adult onset hypertrophic cardiomyopathy in zebrafish. Nature Communications 9 (2018). 10.1038/s41467-018-07054-8

23. Susaki, E. A. et al. Advanced CUBIC protocols for whole-brain and whole-body clearing and imaging. Nature Protocols 10, 1709–1727 (2015). 10.1038/nprot.2015.085

24. Berg, S. et al. ilastik: interactive machine learning for (bio)image analysis. Nat Methods 16, 1226–1232 (2019). 10.1038/s41592-019-0582-9

25. Dobin, A. et al. STAR: ultrafast universal RNA-seq aligner. Bioinformatics 29, 15–21 (2013). 10.1093/bioinformatics/bts635

26. Love, M. I., Huber, W. & Anders, S. Moderated estimation of fold change and dispersion for RNA-seq data with DESeq2. Genome Biology 15 (2014). 10.1186/s13059-014-0550-8

27. Choi, H. M. T. et al. Third-generation in situ hybridization chain reaction: multiplexed, quantitative, sensitive, versatile, robust. Development 145 (2018). 10.1242/dev.165753

28. O’Brown, N. M. et al. The secreted neuronal signal Spock1 promotes blood-brain barrier development. Developmental Cell 58, 1534–1547.e1536 (2023). 10.1016/j.devcel.2023.06.005

